# Embodiment is related to better performance on an immersive brain computer interface in head-mounted virtual reality: A pilot study

**DOI:** 10.1101/578682

**Authors:** Julia M Juliano, Ryan P Spicer, Athanasios Vourvopoulos, Stephanie Lefebvre, Kay Jann, Tyler Ard, Emiliano Santarnecchi, David M Krum, Sook-Lei Liew

**Author notes:** **Correspondence:** Sook-Lei Liew, PhD, OTR/L, University of Southern California, 2025 Zonal Ave., Los Angeles, CA 90033.

## Abstract

Brain computer interfaces (BCI) can be used to provide individuals with neurofeedback of their own brain activity and train them to learn how to control their brain activity. Neurofeedback-based BCIs used for motor rehabilitation aim to ‘close the loop’ between attempted motor commands and sensory feedback by providing supplemental sensory information when individuals successfully establish specific brain patterns. Existing neurofeedback-based BCIs have used a variety of displays to provide feedback, ranging from devices that provide a more immersive and compelling experience (e.g., head-mounted virtual reality (HMD-VR) or CAVE systems) to devices that are considered less immersive (e.g., computer screens). However, it is not clear whether more immersive systems (i.e., HMD-VR) improve neurofeedback performance compared to computer screens, and whether there are individual performance differences in HMD-VR versus screen-based neurofeedback. In this pilot experiment, we compared neurofeedback performance in HMD-VR versus on a computer screen in twelve healthy individuals. We also examined whether individual differences in presence or embodiment correlated with neurofeedback performance in either environment. Participants were asked to control a virtual right arm by imagining right hand movements. Real-time brain activity indicating motor imagery, which was measured via electroencephalography (EEG) as desynchronized sensorimotor rhythms (SMR; 8-24 Hz) in the left motor cortex, drove the movement of the virtual arm towards (increased SMR desynchronization) or away from (decreased SMR desynchronization) targets. Participants performed two blocks of 30 trials, one for each condition (Screen, HMD-VR), with the order of conditions counterbalanced across participants. After completing each block, participants were asked questions relating to their sense of presence and embodiment in each environment. We found that, while participants’ performance on the neurofeedback-based BCI task was similar between conditions, the participants’ reported levels of embodiment was significantly different between conditions. Specifically, participants experienced higher levels of embodiment in HMD-VR compared to the computer screen. We further found that reported levels of embodiment positively correlated with neurofeedback performance only in the HMD-VR condition. Overall, these preliminary results suggest that embodiment may improve performance on a neurofeedback-based BCI and that HMD-VR may increase embodiment during a neurofeedback-based BCI task compared to a standard computer screen.

## 1 Introduction

Neurofeedback training produces beneficial changes in motor function and has been shown to be successful in motor rehabilitation for clinical populations, such as individuals with stroke (Ramos-Murguialday et al., 2013). Neurofeedback-based brain computer interfaces (BCI) use sensory feedback to reward specific patterns of activity in the brain (e.g., as measured with electroencephalography (EEG)). This feedback is then used to control a robotic or computerized device (e.g., movement of an object on a computer screen) to train individuals to control their own brain activity. BCIs designed for the rehabilitation of individuals with severe motor impairment attempt to ‘close the loop’ between motor commands and sensory feedback by providing supplemental sensory information when individuals successfully establish specific brain patterns.

Given that individuals with severe motor impairment cannot generate active volitional movement, a primary neurofeedback approach is to use imagined movement (i.e., motor imagery) to drive the BCI. Motor imagery (MI) is thought to engage areas that modulate movement execution (Dechent et al., 2004; Jackson et al., 2003; Naito et al., 2002). MI has been shown to be an effective intervention for motor rehabilitation, especially when it is coupled with physical practice (Carrasco and Cantalapiedra, 2016; Guerra et al., 2018). Another related approach is to use action observation. The action observation network (AON) consists of motor-related regions in the brain that are active during both the performance of an action and, more importantly, simply during the observation of an action. In this way, action observation provides a feasible way to stimulate cortical motor regions in the absence of volitional movement (Garrison et al., 2010, 2013). Related, action observation therapy, in which patients observe actions that correspond to their paretic limb, has been shown to improve motor rehabilitation in individuals with severe motor impairments (Franceschini et al., 2012; Zhu et al., 2015).

Previous work has shown that neurofeedback-based BCIs employing MI can produce clinically meaningful improvements in motor function in individuals with motor impairments (Ang et al., 2014; Biasiucci et al., 2018; Cincotti et al., 2012; Frolov et al., 2017; Pichiorri et al., 2015; Tung et al., 2013). These neurofeedback-based BCIs have used a variety of displays to provide feedback, ranging from devices that provide an immersive and compelling experience (e.g., projected limbs, robotic orthoses, or exoskeletons; Cincotti et al., 2012; Frolov et al., 2017; Pichiorri et al., 2015; Ramos-Murguialday et al., 2013) to devices that are considered less immersive (e.g., computer screens; Ang et al., 2014; Tung et al., 2013). Recently, BCIs have also begun to incorporate head-mounted virtual reality (HMD-VR) in order to provide a more immersive and realistic environment (Mcmahon and Schukat, 2018) and to provide more biologically relevant feedback (Vourvopoulos and Bermúdez i Badia, 2016). However, it is not known whether HMD-VR improves neurofeedback performance compared to feedback provided on a screen. It is also unclear whether neurofeedback provided in HMD-VR increases one’s feeling of presence and embodiment compared to screen-based neurofeedback.

Studies have shown that HMD-VR facilitates the embodiment of a virtual body and that the observation of this virtual body in the first person perspective is enough to induce a strong feeling of embodiment for the virtual body’s actions (Banakou et al., 2013; Kilteni et al., 2012, 2013; Osimo et al., 2015; Yee and Bailenson, 2007). In HMD-VR, individuals exhibit behaviors that match those of a digital self-representation, such as overestimating object sizes when an adult has been given a virtual child body (Banakou et al., 2013) or exhibiting a reduction in implicit racial bias when given a body of a different race (Banakou et al., 2016). Initially coined the Proteus Effect (Yee and Bailenson, 2007), this sense of embodiment that arises from viewing a virtual limb has the potential to alter one’s own neurophysiology and behavior. Regarding motor behavior, an increased level of presence and embodiment has been shown to be related to increased sensorimotor rhythms (SMR) desynchronization (Vecchiato et al., 2015). Related, observing the actions of virtual limbs in virtual reality have been shown to increase SMR desynchronization (Pavone et al., 2016).

We have created a hybrid brain computer interface for individuals with severe motor impairments called REINVENT (Rehabilitation Environment using the Integration of Neuromuscular-based Virtual Enhancements for Neural Training), which can take brain (EEG) and/or muscle (electromyography (EMG)) signals indicating an attempt to move and provide neurofeedback of an individual’s virtual arm moving in head-mounted virtual reality (HMD-VR). In this way, elements of motor imagery, action observation, and neurofeedback are combined in one platform.

Although we designed REINVENT as a neurofeedback-based BCI device for individuals with severe motor impairments, in this pilot study, we first wanted to examine whether providing neurofeedback in HMD-VR improves neurofeedback performance compared to receiving the same neurofeedback on a computer screen in healthy adults. Furthermore, we wanted to examine whether there were differences in the levels of presence and embodiment induced by HMD-VR versus a computer screen, and how individual differences in these features relate to neurofeedback performance in each environment. As presence and embodiment play an important role in increasing SMR desynchronization and HMD-VR induces high levels of presence and embodiment, we predicted that participants would show better neurofeedback performance in an HMD-VR environment compared to a computer screen, and that improved performance would be related to increased presence and embodiment.

## 2 Materials and Methods

### 2.1 Participants

Twelve healthy participants were recruited for this experiment (7 females/ 5 males; age: M = 24.4 years, SD = 2.7 years). Eligibility criteria included healthy, right handed individuals and informed consent was obtained from all participants. Eight participants reported being naive to head mounted virtual reality; the four participants with previous use of head mounted virtual reality reported using the device no more than four times. The experimental protocol was approved by the University of Southern California Health Sciences Campus Institutional Review Board and performed in accordance with the 1964 Declaration of Helsinki.

### 2.2 REINVENT hardware, software, online processing, and data integration

The REINVENT system is described in technical detail in a previously published paper (Spicer et al., 2017). Briefly, REINVENT (Figure 1A) is a brain computer interface (BCI) that is composed of four main components: electroencephalography (EEG), electromyography (EMG), an inertial measurement unit (IMU), and a head-mounted virtual reality (HMD-VR) system. Custom software is used to control the BCI and provide users with real-time feedback of a virtual arm. EEG signals were recorded from electrodes of interest over the left motor cortex (i.e., C1, C3, and CP1, based on the international 10-20 system) with both ear lobes used as the ground and reference electrodes, and sent to the REINVENT software. Data processing occurred online. Individual channels were high-pass filtered using a second order Butterworth filter with a cutoff of 3 Hz, and a sliding window consisting of 125 incoming samples were fast Fourier transformed (FFT). Power was then computed between the frequency ranges of 8-24 Hz, capturing the broad activity in alpha and beta bands that may correspond to motor imagery (i.e., sensorimotor desynchronization). The virtual arm direction updated every second and moved towards the target in response to sensorimotor desynchronization, measured as a decrease in amplitude compared to the baseline recording of the left sensorimotor area (i.e., the combined three channels: C1, C3, CP1).

**Figure 1.**
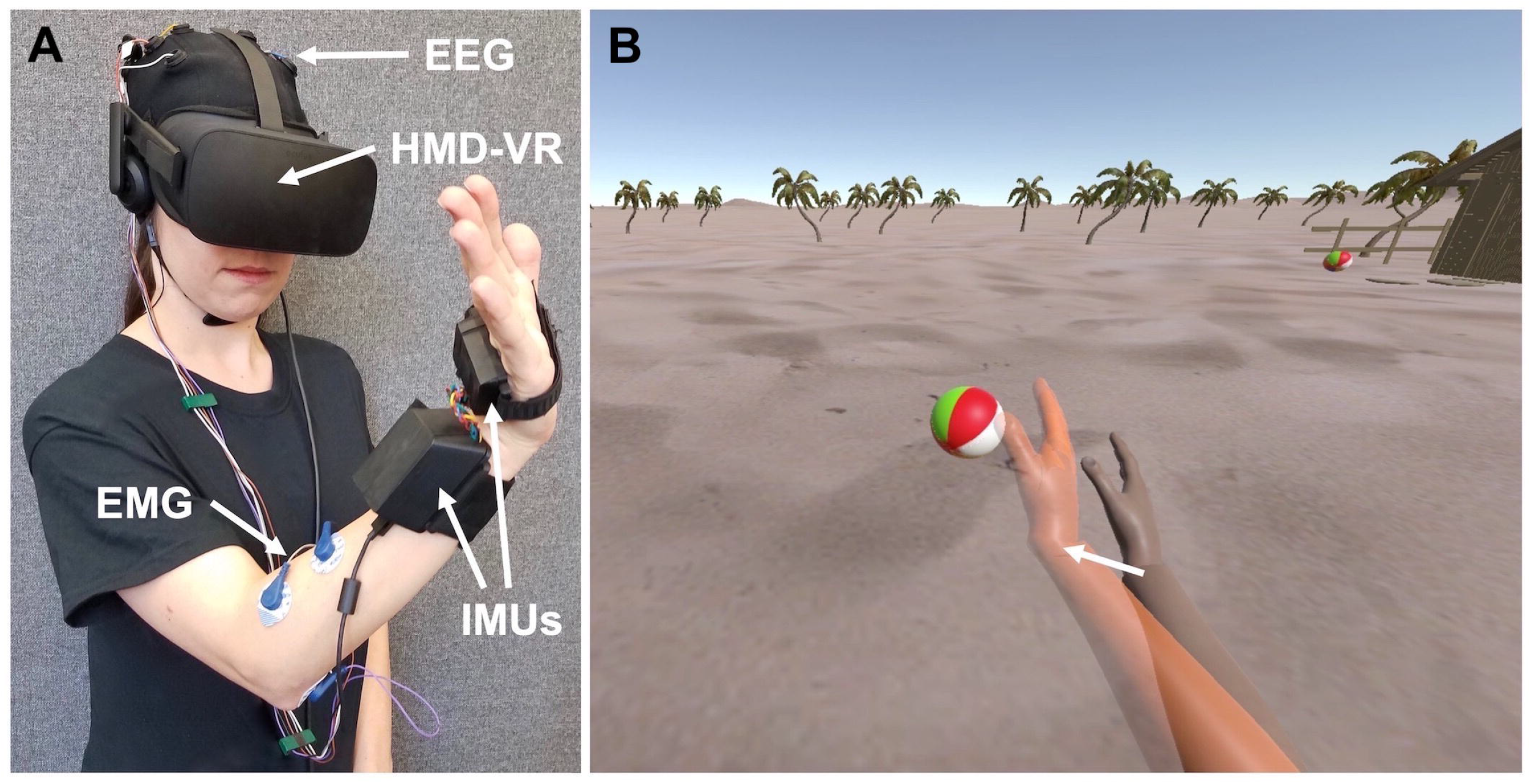
REINVENT system. **(A)** REINVENT hardware used here is composed of electroencephalography (EEG), electromyography (EMG), inertial measurement units (IMUs), and a head-mounted virtual reality (HMD-VR) system. Written informed consent for the publication of this image was obtained from the individual depicted. **(B)** The environment participants observed on both a computer screen and in HMD-VR; arm movements are goal-oriented such that when the arm reaches a target position, it interacts with an object (e.g., hitting a beach ball). On EEG blocks (Screen, HMD-VR), participants would attempt to move their virtual arm (right arm) to the orange target arm (left arm) by thinking about movement. On the IMU block, the virtual arm would match participants actual arm movements.

#### 2.2.1 Electroencephalography (EEG) and electromyography (EMG)

The EEG/EMG component of REINVENT is composed of hardware from OpenBCI (www.openbci.com), a low-cost solution for measuring brain and muscle activity. The EEG component consists of reusable dry EEG electrodes and the EMG component consists of snap electrode cables connected to mini disposable gel electrodes (Davis Medical Electronics, Inc.). Both EEG and EMG wires were connected to a 16-channel, 32-bit v3 processor (Cyton + Daisy Biosensing OpenBCI Board) and sampled at 125 Hz.

Twelve EEG locations based on the international 10-20 system and concentrated over the prefrontal and motor cortex were used to record brain activity (F3, F4, C1, C2, C3, C4, CP1, CP2, CP5, CP6, P3, and P4). Ground and reference electrodes were located at the right and left earlobes, respectively. For the neurofeedback, the sum desynchronization from C1, C3 and CP1, representing the left motor network, was used to drive the movement of a virtual right arm towards a target arm. EMG was recorded from four electrodes placed on the wrist flexors and extensors on the muscle bellies of the right forearm, with a reference electrode on the bony prominence of the elbow. In the current experiment, muscle activity from EMG was collected but not analyzed or reported.

#### 2.2.2 Arm movement

To foster a sense of embodiment between the participant and the virtual arm, the participant’s own arm movements were recorded using two Nine Degrees of Freedom (9DOF) IMUs, with one placed on the hand and the other placed on the wrist of the right arm (Spicer et al., 2017). Before beginning the experiment, the participant’s arm was passively moved by the experimenter and the virtual representation of the arm was shown on the computer screen and in HMD-VR. In this way, a sensorimotor contingency was developed between the participant’s own arm and the virtual arm they were subsequently asked to control.

### 2.3 Displays

For the HMD-VR condition, we used the Oculus CV1 which includes positional and rotational tracking to display the stimuli. For the Screen condition, we used a 24.1 inch, 1920 × 1200 pixel resolution computer monitor (Hewlett-Packard) to display the stimuli. In both displays, participants observed a scene that included two virtual arms: (1) one virtual arm that represented the participant’s own arm and (2) a second virtual arm, colored in orange, that provided different target arm positions that participants were asked to move their own arm towards (Figure 1B).

### 2.4 Experimental design

All participants underwent the same experimental design and completed all conditions (Screen, HMD-VR). Prior to the experiment, a resting EEG baseline of three minutes with the HMD-VR removed was recorded for each participant. Participants were instructed to keep their eyes open and fixed on a location at the center of the computer screen. For the duration of the recording, participants were asked to think about a stationary object and to stay as still as possible. The recording was used to provide the baseline EEG values for the experiment. Participants then completed three blocks of 30 trials (90 trials in total) where each block was a separate condition. The conditions were (1) controlling the virtual arm with brain activity on the computer screen (Screen), (2) controlling the virtual arm with brain activity in head-mounted virtual reality (HMD-VR), and (3) controlling the virtual arm with actual arm movements in head-mounted virtual reality (IMU). Participants completed the conditions in the following block order: Block 1 (Screen), Block 2 (HMD-VR), Block 3 (IMU), with the first two blocks (Screen, HMD-VR) counterbalanced. In this experiment, the IMU condition strictly provided a control condition of real movement instead of neurofeedback; this data is briefly reported but not focused on in this paper. Before starting the experimental conditions, participants were given instructions on how to control their virtual arm (i.e., “You will see two right arms. One is orange and that is the target arm that moves to different positions. The other is your arm. We want you to move it to match the target arm’s position. You can move your arm in two ways. First, you will complete 60 trials of moving the virtual arm with just your thoughts by thinking about moving; 30 of the trials will be on the computer screen, without the head-mounted virtual reality, and 30 trials will be with the head-mounted virtual reality. Then you will complete 30 trials of moving the virtual arm using your actual arm movements.”). Instructions were repeated at the start of each condition. After the completion of each EEG neurofeedback condition (Screen, HMD-VR), a resting-EEG acquisition of three minutes was recorded while the HMD-VR was removed; participants were again instructed to keep their eyes open and fixed on the center of the screen for the duration of the recording. Figure 2 shows a detailed timeline of the experimental design.

**Figure 2.**
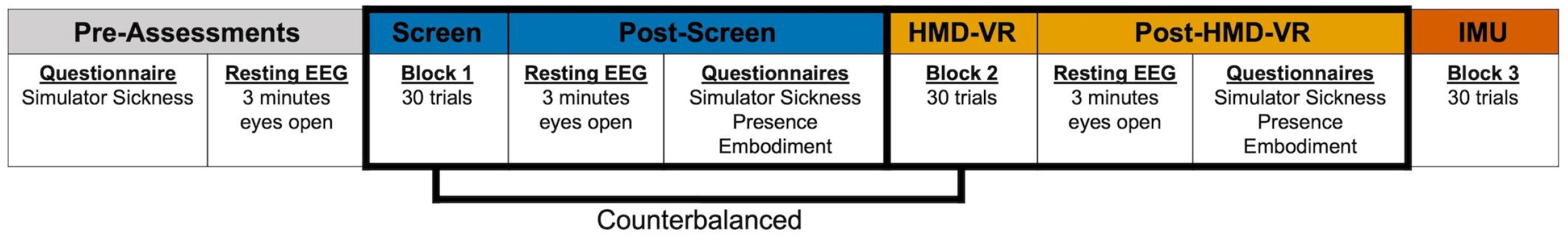
Experimental timeline. Prior to the experimental blocks, participants completed a questionnaire relating to simulator sickness and then completed a resting EEG recording for three minutes with eyes open. Participants then completed the three experimental blocks where the first two blocks were counterbalanced; during Blocks 1 and 2 (Screen, HMD-VR), participants were asked to think about movement in order to move their virtual arm to a virtual target arm on either a computer screen or in HMD-VR. After the Screen condition and after the HMD-VR condition, participants completed a resting EEG recording for three minutes with eyes open and then completed a series of questionnaires relating to simulator sickness, presence, and embodiment. During Block 3 (IMU), participants were asked to move their physical arm to a virtual target arm in HMD-VR.

#### 2.4.1 Individual trials

At the start of each trial, a target arm animated a wrist extension pose in one of three target positions. Once the target arm stopped moving, participants were instructed to move their virtual arm to match the position of the target arm given the current condition (i.e., in the case of the two EEG neurofeedback conditions (Screen, HMD-VR), they were asked to think about moving; in the case of the IMU condition, they were asked to actually move their arm to the target location). During the EEG neurofeedback condition trials, the virtual hand incremented either forward or backward, as determined by the sum of the three channel EEG desynchronization compared to baseline. Most of the time, the EEG activity was significantly above or below the baseline; however, if the sensorimotor activity was hovering around the baseline, the arm would move back and forth. The duration of each trial was 15 seconds; if the target arm was reached within this time constraint, a successful auditory tone was played, however, if the target arm was not reached, then an unsuccessful auditory tone was played. At the completion of each trial, the target and virtual arms returned to their starting position.

### 2.5 Subjective Questionnaires

Prior to the experiment, participants were given a series of standard questions about their baseline comfort levels (Simulator Sickness Questionnaire; adapted from Kennedy et al. 1993). After participants completed each EEG neurofeedback condition (Screen, HMD-VR), they were given the same simulator sickness questionnaire to examine changes following each block. Responses were reported on a 0 to 3-point scale and questions were collapsed along three main features: Nausea, Oculomotor, and Disorientation. In addition, after completing both the Screen and HMD-VR conditions, participants were also asked questions pertaining to their overall sense of presence and embodiment in each respective environment. The Presence Questionnaire was adapted from Witmer and Singer (1998) and revised by the UQO Cyberpsychology Lab (2004) and asked participants a series of questions to gauge their sense of presence in each environment. Responses were reported on a 1 to 7-point scale and questions were collapsed along five main features: Realism, Possibility to Act, Quality of Interface, Possibility to Examine, and Self-Evaluation of Performance. The Embodiment Questionnaire was adapted from Bailey et al. (2016) and Banakou et al. (2013) and asked participants a series of questions to gauge their sense of embodiment. Responses were reported on a 1 to 10-point scale and questions were averaged to generate an overall Embodiment feature. In addition, we also collapsed questions relating to either Self Embodiment or Spatial Embodiment to generate two embodiment sub-features. Table 1 includes individual questions asked on the Embodiment Questionnaire.

**Table 1.** Individual Questions on Embodiment Questionnaire. After the Screen and HMD-VR conditions (Blocks 1, 2), participants were asked questions relating to their level of embodiment in each of the respective conditions. Participants reported their level of embodiment on a scale from 1 to 10. Self Embodiment and Spatial Embodiment was calculated by averaging the responses given for each respective question type.

| Type | Question | Referenced | Scoring Scale |
| --- | --- | --- | --- |
| Self | To what extent did you feel that the virtual arm was your own arm? | Own Arm | Not at all/Very much (1...10) |
| Self | How much did the virtual arm's actions correspond with your commands? | Arms Actions | Not at all/Very much (1...10) |
| Self | To what extent did you feel if something happened to the virtual arm it felt like it was happening to you? | Happening to Arm | Not at all/Very much (1...10) |
| Self | How much control did you feel you had over the virtual arm in this virtual environment? | Amount of Arm Control | No control/Full control (1...10) |
| Self | How much did you feel that your virtual arm resembled your own (real) arm in terms of shape, skin tone or other visual features? | Resembled Arm | Not at all/Very much (1...10) |
| Self | Did the virtual arm seem bigger, smaller or about the same as what you would expect from your everyday experience? | Size of Arm | Smaller/Larger (1...10) |
| Spatial | To what extent did you feel like you were really located in the virtual environment? | Location | None/Completely (1...10) |
| Spatial | To what extent did you feel surrounded by the virtual environment? | Surrounded | None/Completely (1...10) |
| Spatial | To what extent did you feel that the virtual environment seemed like the real world? | Real World | None/Completely (1...10) |
| Spatial | To what extent did you feel like you could reach out and touch the objects in the virtual environment? | Reach Out and Touch | None/Completely (1...10) |

### 2.6 Analyses

#### 2.6.1 Post-hoc EEG analysis on activity during task

In addition to the online processing (see section 2.2), post-hoc EEG signals were processed offline using MATLAB^®^ (R2017a, The MathWorks, MA, USA) with the EEGLAB toolbox (Delorme and Makeig, 2004). After importing the data and channel information, a high-pass filter at 1 Hz was applied to remove the ‘baseline drift’ followed by line-noise and harmonics removal at 60 Hz. Furthermore, bad channels were rejected while any potential missing channels were interpolated before the re-referencing stage. Additionally, all channels were re-referenced to the average. Next, data epoching was performed by extracting the trials from the EEG neurofeedback conditions (Screen, HMD-VR) for each participant. Finally, the baseline data (180 seconds) were extracted from the resting-state session that occurred before the task.

For computing the average spectral power, Welch’s method for Power Spectral Density (PSD) of the power spectrum (Welch, 1967) was used across the online frequency range (8-24 Hz) and for the alpha (8-12 Hz) and beta (13-24 Hz) bands. PSD was extracted from both the epoched motor-related data and the baseline. Finally, the band power was extracted over the C3 electrode location and calculated using the following formula:

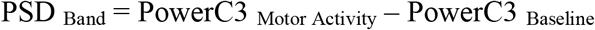

#### 2.6.2 Statistical Analysis

Statistical analysis for neurofeedback performance, subjective experience from questionnaires, and EEG activity during the task was analyzed using the statistical package R (3.2.2, The R Foundation for Statistical Computing, Vienna, Austria). To assess statistical differences in performance, subjective experience, and average spectral power during the task between the two EEG conditions (Screen, HMD-VR), a paired t-test was performed on each measure. Means (M) and standard deviations (SD) are reported for each measure. To confirm that neurofeedback based on motor imagery was successfully used to increase performance, we ran a simple linear regression on neurofeedback performance based on PSD. Lastly, we examined the relationship between neurofeedback performance and responses from the Presence Questionnaire and the Embodiment Questionnaire using regression analysis. For the Presence Questionnaire, we ran a multiple regression analysis on neurofeedback performance based on the five presence features for each condition (Screen, HMD-VR). For the Embodiment Questionnaire, we first ran a simple linear regression analysis on neurofeedback performance based on the overall Embodiment feature for each condition. Then, we ran a multiple regression analysis on neurofeedback performance based on the two embodiment sub-features (Self Embodiment and Spatial Embodiment) for each condition. For all regression analyses, adjusted R^2^ is reported. All participants completed the IMU condition with 100% accuracy and therefore this condition is not included in this analysis.

## 3 Results

### 3.1 Differences in neurofeedback performance and time to complete trials between Screen and HMD-VR

The proportion of correct trials completed was similar between the two conditions (Figure 3A; t(11) = −0.46, p = 0.656; Screen: M = 80.95%, SD = 9.1%, and HMD-VR: M = 83.33%, SD = 14.9%). These results suggest that participants seemed to perform similarly independent of whether neurofeedback was provided in HMD-VR or on a computer screen.

**Figure 3.**
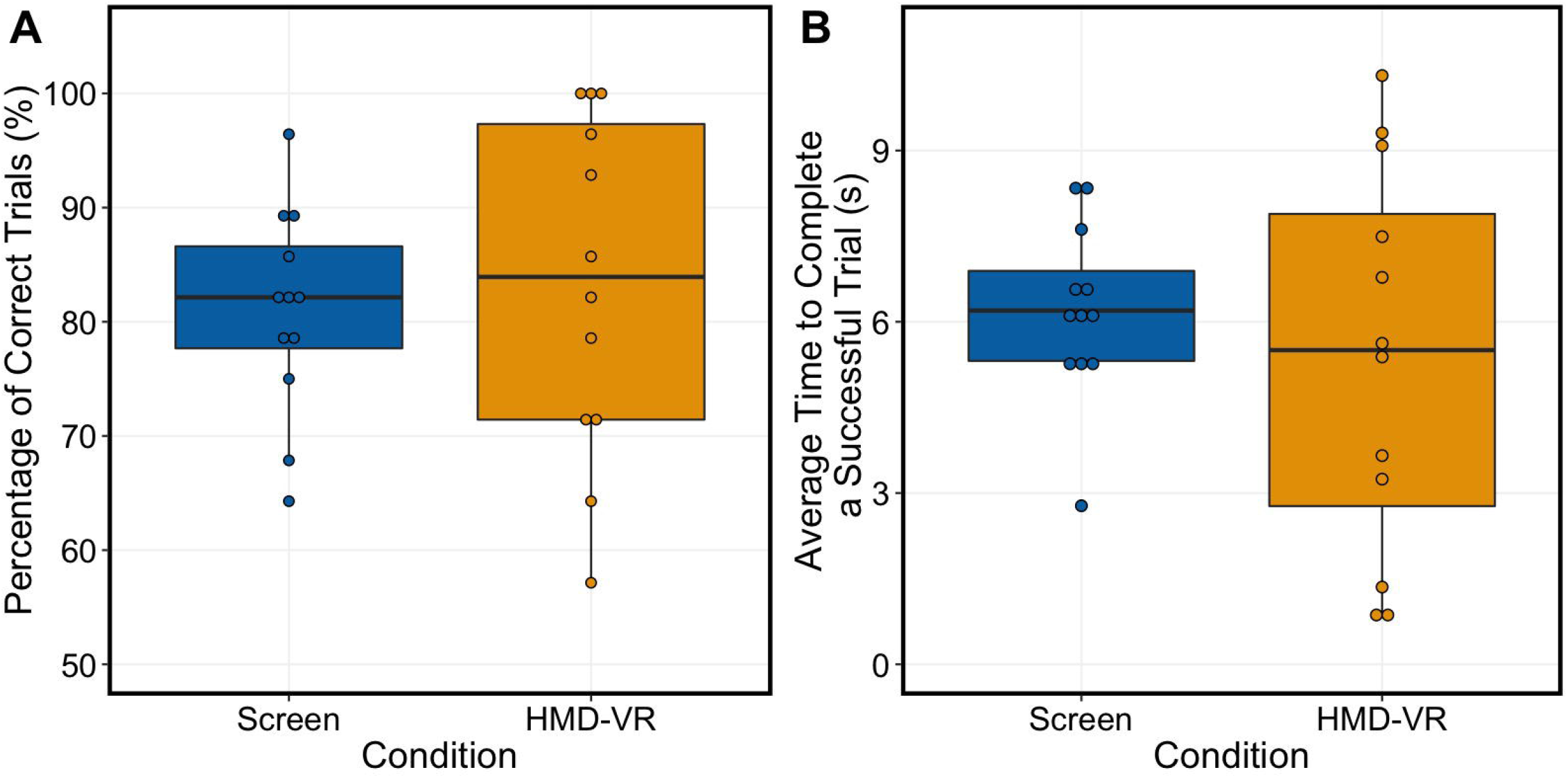
Average performance on trials and time to complete successful trials between conditions. **(A)** The analysis showed no significant differences in performance between Screen (left, blue) and HMD-VR (right, yellow) conditions (t(11) = −0.46, p = 0.656). **(B)** The analysis showed no significant differences in time on successful trials between Screen (left, blue) and HMD-VR (right, yellow) conditions (t(11) = 0.54, p = 0.597).

Similarly, the time to complete each of the successful trials was also similar between the two conditions (Figure 3B; t(11) = 0.54, p = 0.597; Screen: M = 4.347 s, SD = 1.17 s, and HMD-VR: M = 3.996 s, SD = 2.41 s). These results suggest that when participants were able to increment the virtual arm towards the target with their brain activity, the efficiency of control was similar whether viewing the arm in the HMD-VR environment or on a computer screen.

### 3.2 Differences in power spectral density between Screen and HMD-VR

Similar to the neurofeedback performance results, we did not find significant differences in group-level PSD between the Screen and HMD-VR conditions across the 8-24 Hz frequency range (Figure 4A; t(11) = 0.475, p = 0.644; Screen: M = −4.69, SD = 2.96, and HMD-VR: M = −4.32, SD = 3.41). We also explored alpha and beta bands separately, and did not find significant differences in group-level PSD between the Screen and HMD-VR conditions in either band (alpha: Figure 4B, t(11) = 1.363, p = 0.200, Screen: M = −1.84, SD = 2.90, and HMD-VR: M = −2.89, SD = 3.04; beta: Figure 4C, t(11) = −1.141, p = 0.278, Screen: M = −5.88, SD = 3.08, and HMD-VR: M = −4.92, SD = 3.63). This further suggests that participants had similar levels of sensorimotor activity whether neurofeedback was provided in HMD-VR or on a computer screen. Additionally, we have included two supplementary figures reporting individual participant EEG activity in alpha and beta bands for both C3 (Supplementary Figure 1; contralateral to and controlling of the virtual hand) and C4 (Supplementary Figure 2; ipsilateral to the virtual hand) recordings.

**Figure 4.**
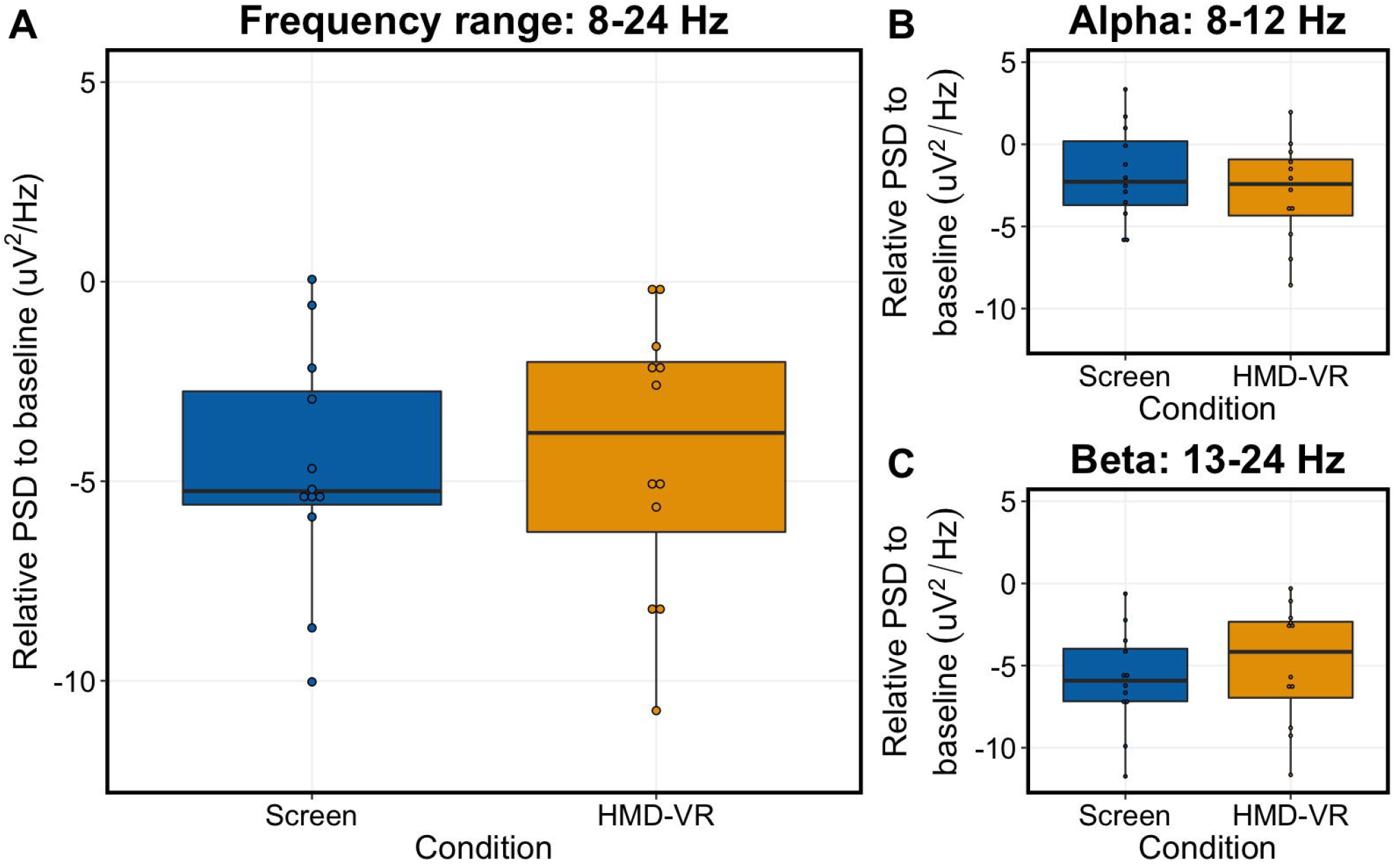
Average power spectral density during trials between conditions. **(A)** The relative group-level PSD (8-24 Hz) between the Screen (left, blue) and HMD-VR (right, yellow) conditions was not significantly different (t(11) = 0.475, p = 0.644). **(B)** The relative group-level alpha between the Screen (left, blue) and HMD-VR (right, yellow) conditions was also not significantly different (t(11) = 1.363, p = 0.200). **(C)** The relative group-level beta between the Screen (left, blue) and HMD-VR (right, yellow) conditions was also not significantly different (t(11) = −1.141, p = 0.278).

### 3.3 Relationship between power spectral density and neurofeedback performance

To confirm the relationship between PSD in the 8-24 Hz frequency range and the corresponding neurofeedback performance, we ran a simple linear regression of neurofeedback performance based on PSD. As expected, we found a significant relationship between PSD and neurofeedback performance (Figure 5; F(1,22) = 9.328, p = 0.006; R^2^ = 0.266) where an increased sensorimotor desynchronization corresponded to better neurofeedback performance.

**Figure 5.**
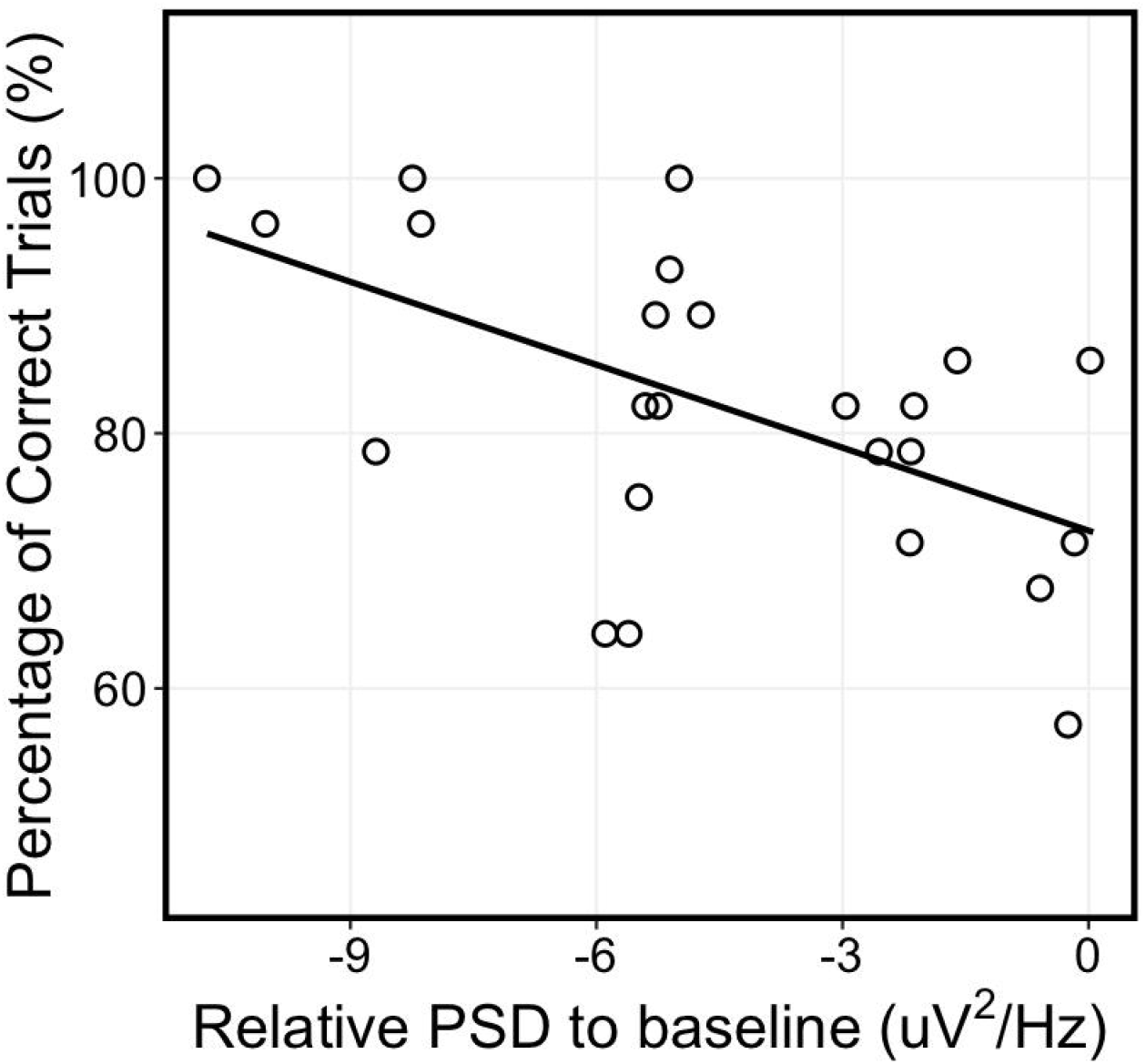
Relationship between power spectral density and neurofeedback performance. There was a significant relationship between PSD and neurofeedback performance (F(1,22) = 9.328, p = 0.006; R^2^ = 0.266).

### 3.4 Differences in subjective experience between Screen and HMD-VR

There was a significant difference in reports of embodiment between the two conditions (t(11) = - 2.21, p = 0.049; Screen: M = 4.68, SD = 1.27, and HMD-VR: M = 5.4, SD = 1.71) where individuals reported higher levels of Embodiment in the HMD-VR condition. We then examined the sub-features of embodiment, and found a significant difference in reports of spatial embodiment between the two conditions (t(11) = −3.77, p = 0.003; Screen: M = 3.60, SD = 2.04, and HMD-VR: M = 5.35, SD = 2.00) where individuals reported higher levels of Spatial Embodiment in the HMD-VR condition. However, there was no significant difference in reports of self embodiment between the two conditions (t(11) = −0.10, p = 0.922; Screen: M = 5.39, SD = 1.17, and HMD-VR: M = 5.43, SD = 1.76). These results suggest that neurofeedback presented in HMD-VR increases one’s feeling of embodiment compared to neurofeedback presented on a computer screen.

In contrast, there were no significant differences in reports of simulator sickness between the Screen (Nausea: M = 1.59, SD = 8.94; Oculomotor: M = 9.48, SD = 12.15; Disorientation: M = 4.64, SD = 17.13) and the HMD-VR (Nausea: M = 2.39, SD = 5.93; Oculomotor: M = 9.45, SD = 9.76; Disorientation: M = 3.48, SD = 8.65) conditions (Nausea: t(11) = −0.56, p = 0.586; Oculomotor: t(11) = 0.00, p = 1.00; Disorientation: t(11) = 0.43, p = 0.674). These results suggest that HMD-VR neurofeedback does not cause additional adverse effects beyond using a computer screen in healthy individuals.

In addition, there were no significant differences between reports of presence in the two conditions (Realism: t(11) = −1.95, p = 0.078, Screen: M = 30.00, SD = 6.35, HMD-VR: M = 33.00, SD = 6.40; Possibility to Act: t(11) = −1.37, p = 0.199, Screen: M = 18.17, SD = 3.70, HMD-VR: M = 19.92, SD = 4.19; Quality of Interface: t(11) = – 0.62, p = 0.548, Screen: M = 12.83, SD = 3.07, HMD-VR: M = 13.42, SD = 2.97; Possibility to Examine: t(11) = – 2.01, p = 0.070, Screen: M = 13.17, SD = 2.59, HMD-VR: M = 14.92, SD = 2.27; Self-Evaluation of Performance: t(11) = −1.24, p = 0.241, Screen: M = 10.0, SD = 1.95, HMD-VR: M = 11.00, SD = 2.13). This suggests that HMD-VR neurofeedback may specifically increase embodiment but not presence in healthy individuals.

### 3.5 Relationship between embodiment, presence, and neurofeedback performance

We next examined whether individual differences in embodiment related to neurofeedback performance for each condition. We ran a simple linear regression of neurofeedback performance based on the overall Embodiment feature. For the HMD-VR condition, we found a significant relationship between embodiment and neurofeedback performance (F(1,10) = 8.293, p = 0.016; R^2^ = 0.399). However, for the Screen condition, we did not find a significant relationship between embodiment and neurofeedback performance (F(1,10) = 0.434, p = 0.525; R^2^ = −0.054). These results suggest that level of embodiment is specifically related to neurofeedback performance only in HMD-VR and not on a computer screen (Figure 6A; yellow regression line).

**Figure 6.**
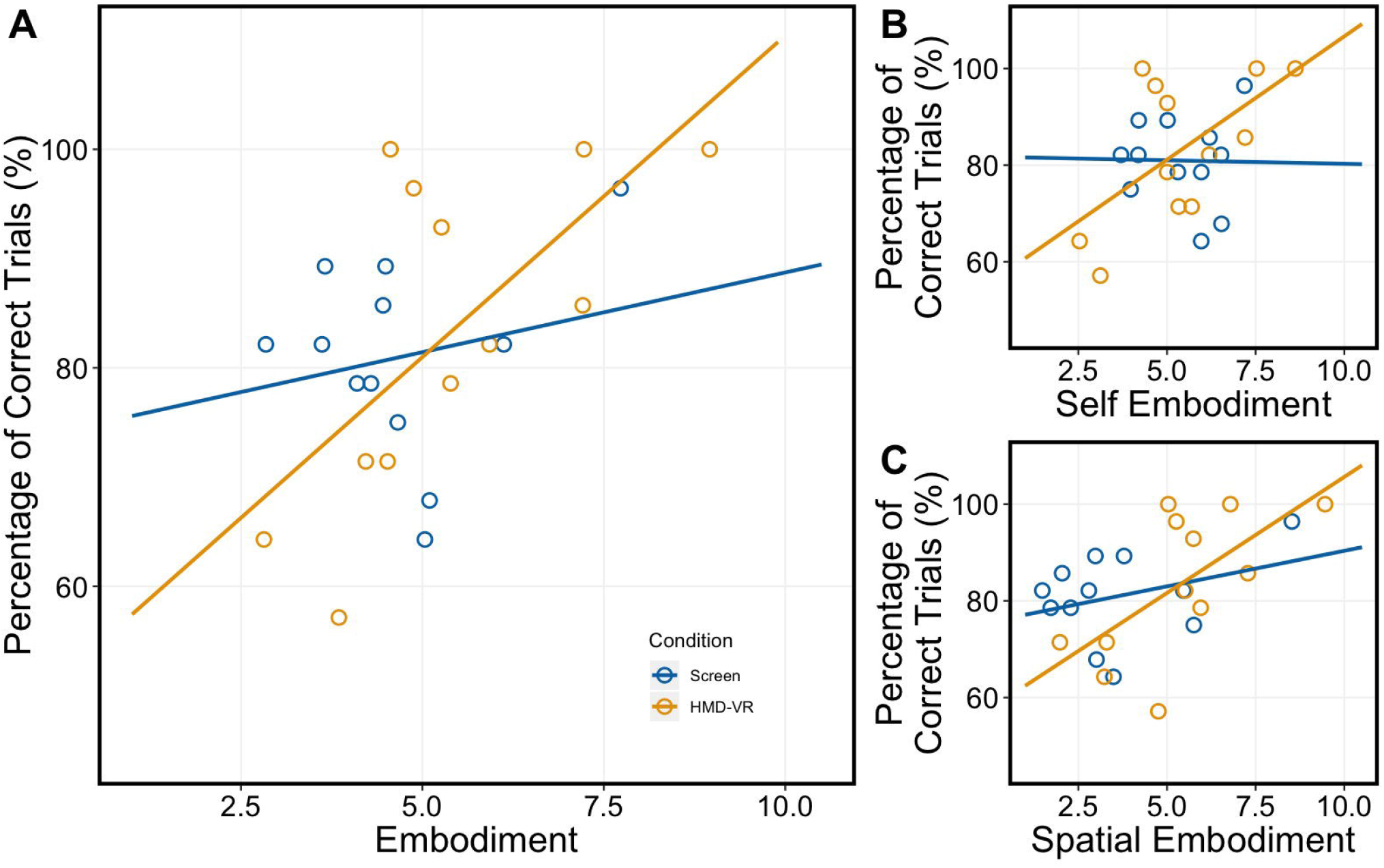
Relationship between subjective experience and neurofeedback performance in Screen and HMD-VR. Participants reported their level of Embodiment on a scale from 1 to 10 (Table 1). **(A)** Embodiment: For the HMD-VR condition, embodiment was significantly related to performance (F(1,10) = 8.293, p = 0.016; R^2^ = 0.399; yellow). However, for the Screen condition, embodiment did not significantly relate to neurofeedback performance (F(1,10) = 0.434, p = 0.525; R^2^ = −0.054; blue). **(B)** Self Embodiment and **(C)** Spatial Embodiment: For the HMD-VR condition, we found a near significant relationship between the two embodiment sub-features and neurofeedback performance (F(2,9) = 3.858, p = 0.0617; R^2^ = 0.342; yellow). However, for the Screen condition, we did not find a significant relationship between the two embodiment sub-features and neurofeedback performance (F(2,9) = 0.706, p = 0.519; R^2^ = −0.056; blue).

To better understand whether specific sub-features of embodiment also related to neurofeedback performance, we then examined if participants’ levels of self and spatial embodiment related to their neurofeedback performance for each condition (Screen, HMD-VR). We ran a multiple linear regression of neurofeedback performance based on the two embodiment sub-features (i.e., Self Embodiment, Spatial Embodiment). For the HMD-VR condition, we found a near significant relationship between the two embodiment sub-features and neurofeedback performance (F(2,9) = 3.858, p = 0.0617; R^2^ = 0.342). For the Screen condition, we did not find a significant relationship between the two embodiment sub-features and neurofeedback performance (F(2,9) = 0.706, p = 0.519; R^2^ = −0.056). These results further suggest that level of embodiment is specifically related to HMD-VR neurofeedback performance. Figure 6B and 6C show regression lines for both Self Embodiment and Spatial Embodiment, respectively.

Although there were no differences in presence between the Screen and HMD-VR conditions, we also explored whether individual differences in presence related to neurofeedback performance for each condition (Screen, HMD-VR). We ran a multiple linear regression of neurofeedback performance based on the five presence features (i.e., Realism, Possibility to Act, Quality of Interface, Possibility to Examine, and Self-Evaluation of Performance). We did not find a significant relationship between the five presence features and neurofeedback performance for either the Screen or HMD-VR condition (HMD-VR: (F(5,6) = 0.476, p = 0.452; R^2^ = 0.039); Screen: F(5,6) = 0.840, p = 0.567; R^2^ = −0.078). These results suggest that the level of presence does not seem to be significantly related to either HMD-VR or computer screen neurofeedback performance.

## 4 Discussion

The current pilot study examined whether neurofeedback from a motor-related brain computer interface provided in HMD-VR could lead to better neurofeedback performance compared to the same feedback provided on a standard computer screen. In addition, differences in embodiment and presence between Screen and HMD-VR conditions were examined. Finally, we explored whether individual differences in embodiment and presence related to neurofeedback performance in each condition. Overall, we found preliminary evidence that healthy participants showed similar levels of neurofeedback performance in both Screen and HMD-VR conditions; however, we found a trend for better performance in the HMD-VR condition. Additionally, participants reported greater embodiment in the HMD-VR versus Screen condition, and higher reported levels of embodiment related to better neurofeedback performance in the HMD-VR condition only. These preliminary results suggest that HMD-VR-based neurofeedback may rely on an individual’s sense of embodiment for successful performance. Future studies should explore these findings with a larger sample size over a longer period of time.

### 4.1 Neurofeedback performance between a computer screen and HMD-VR

Regardless of condition (Screen, HMD-VR), we found that on average, individuals were able to accurately modulate their brain activity to successfully control a virtual arm on over 80 percent of trials. These results suggest that neurofeedback based on motor imagery, using biologically-relevant stimuli, can occur either on a computer screen or in head-mounted virtual reality. However, as seen in Figures 3A and 3B, there is a trend towards better performance and faster time to complete a successful trial in the HMD-VR condition compared to the Screen condition, which may not allow for significance because of our limited dataset (further discussed in section 4.6). This trend towards greater sensorimotor desynchronization can also be observed in the individual subject data (Supplementary Figures 1 and 2), with more individuals showing more sensorimotor activity for the HMD-VR condition than the Screen condition. Additionally, there is a larger range of interindividual variability in both performance and average time to complete a successful trial in the HMD-VR condition, suggesting that some individuals may benefit from HMD-VR compared to others. This suggestion is further supported by the correlation between performance and embodiment, in which we show that individuals who had greater embodiment had better performance in HMD-VR only (further discussed in section 4.4).

### 4.2 Power spectral density between a computer screen and HMD-VR

Similarly, regardless of condition (Screen, HMD-VR), we found that on average, individuals had similar levels of sensorimotor activity, as measured by PSD between 8-24 Hz, and when divided into alpha and beta frequency bands. This was expected as the sensorimotor desynchronization used to calculate PSD was also used to drive the virtual arm in the task. However, similar to the performance results, we see a trend for greater desynchronization in the alpha band for the HMD-VR condition (Figure 4B). While we do not see a trend for greater desynchronization in the beta band for the HMD-VR condition (Figure 4C), these results may indicate a neurofeedback-based effect for the different displays, suggesting that feedback type may be able to alter brain activity. We also showed a significant relationship between PSD and neurofeedback performance, where increased desynchronization corresponded to increased performance.

### 4.3 A higher level of embodiment in HMD-VR compared to a computer screen

After performing the neurofeedback task in each condition (Screen, HMD-VR), participants reported having higher levels of embodiment in HMD-VR compared to the computer screen. This is in agreement with previous research showing that HMD-VR is effective for inducing embodiment (Osimo et al., 2015; Slater and Sanchez-Vives, 2016). However, while it has been intuitively suggested that viewing a virtual body in HMD-VR should induce greater embodiment than viewing the same virtual body on a computer screen, to our knowledge, there has been little empirical evidence to demonstrate this. Here, we address this gap by providing evidence that HMD-VR does seem to in fact increase embodiment compared to a computer screen during a neurofeedback task.

### 4.4 Greater embodiment is related to better neurofeedback performance in HMD-VR

In line with our hypothesis, we show that greater embodiment was positively related to better neurofeedback performance in HMD-VR. This uniqueness to HMD-VR could possibly be explained by an increased range of embodiment levels in the HMD-VR condition compared to the Screen condition. These results are consistent with previous research where embodiment has been shown to lead to neurophysiological and behavioral changes based on the virtual body’s characteristics, such as overestimating object distances after given an elongated virtual arm in HMD-VR (Kilteni et al., 2012). While these findings do not support causality, they are important because they suggest that embodiment may have the potential to improve an individual’s neurofeedback performance, and HMD-VR may be able to increase the level of embodiment of an individual, beyond that of a normal computer screen. This suggests that if individuals were to encounter a ceiling effect while controlling neurofeedback on a computer screen, they might be able to show greater improvements, beyond this ceiling, if they show greater embodiment in HMD-VR.

### 4.5 Future clinical implications

We designed REINVENT as a neurofeedback-based BCI for individuals with severe motor impairments, such as stroke. However, before exploring the effectiveness of this device in a population with severe motor impairments, we first examined whether providing neurofeedback in HMD-VR improves performance compared to receiving the same neurofeedback on a computer screen in healthy adults. Our findings suggest that increased embodiment may improve individuals’ neurofeedback performance, which could potentially improve patients’ recovery. Furthermore, our results suggest that HMD-VR may facilitate an increased level of embodiment, beyond what might be seen with traditional screen-based BCIs. As previous brain computer interfaces have been shown to have a positive change on muscle and sensorimotor brain activity in post-stroke individuals, even when using screen-based environments (Ono et al., 2014), we anticipate that embodiment in HMD-VR may lead to even greater improvements. Future work might explore whether additional measures of embodiment, administered prior to HMD-VR neurofeedback training, could predict embodiment and neurofeedback performance. If so, these “pre-assessments” of embodiment potential could be used to predict and personalize neurofeedback-based BCI therapy. However, as this data is preliminary, more data is needed to explore this hypothesis.

### 4.6 Limitations

Our pilot study has several limitations. First was the limited sample size of 12 individuals and the limited number of trials collected per condition (i.e., 30 trials per condition). However, even with this limited sample, we were still able to extract the power spectral density (PSD), calculate relative PSD to baseline, and find a significant relationship between PSD and neurofeedback performance. However, future research should explore this with greater power both in the number of participants and in the number of trials collected.

A second limitation was the lack of longitudinal data collected in the experiment, which limited the potential amount of training participants received and the potential for performance improvement. However, as this was a pilot study, we only collected data during one session. In future studies, we plan to replicate this experiment, including additional trials and sessions in the experimental design.

A third limitation was the use of only 8 channels of dry electrodes to collect sensorimotor activity and the broad frequency band used (8-24 Hz). Given that our system was initially designed to provide a low-cost rehabilitation intervention, we chose to drive the neurofeedback-based device with a limited number of dry electrodes as previous studies have found dry electrodes to be suitable for neurofeedback applications (Mcmahon and Schukat, 2018; Uktveris and Jusas, 2018). However, we recognize that the signal quality of these electrodes can be noisy, and even though we were able to successfully extract power spectral density, in future studies, we plan to use higher quality electrodes (e.g., active gel electrodes) which would also allow us to narrow the frequency band and personalize the feedback across individuals. In addition, although the low resolution from 8 channels, primarily clustered around bilateral sensorimotor regions, facilitated a faster application of the EEG cap, it also limited our post-hoc analyses. Future research studies should utilize more channels for higher resolution. This would enable topographical analyses of whole brain activity during neurofeedback training as well as the ability to examine brain activity in non-motor regions as control regions.

A fourth limitation is that here, we studied only healthy individuals. This is notable as the effects observed may be smaller than those of a clinical population, who may have more room to improve. Specifically, the healthy individuals in our study showed, on average, 80% accuracy with the neurofeedback-based BCI within a short time frame, which may reflect their intact sensorimotor control. However, individuals with severe motor impairments may start with lower scores and have greater room for improvement due to damage to these same networks. Future work may examine extended training with the HMD-VR environment to see if it is possible for individuals to improve beyond their current levels with greater time in the environment, as well as the effects of embodiment on neurofeedback-based BCI performance in individuals with stroke, which may provide a greater range of abilities and thus greater potential effects with immersive virtual reality. Future work should build upon these modest results and explore the effects of embodiment on HMD-VR neurofeedback performance with large samples and in clinical populations.

### 4.7 Conclusions

This preliminary work suggests that individuals have higher levels of embodiment when given immersive virtual reality-based neurofeedback compared to the neurofeedback displayed on a computer screen. Furthermore, this increased sense of embodiment in immersive virtual reality neurofeedback has the potential to improve neurofeedback performance in healthy individuals over their performance on a computer screen. HMD-VR may provide a unique medium for improving neurofeedback-based BCI performance, especially in clinical settings related to motor recovery. Future work will explore ways to increases presence and embodiment in immersive head-mounted virtual reality and examine these effects on motor rehabilitation in patients with severe motor impairment.

## 5 Acknowledgments

We thank David Saldana for assistance with recruitment and Catherine Finnegan for assistance with recruitment and initial pilot data collection.

## Reference

Ang, K. K., Guan, C., Phua, K. S., Wang, C., Zhou, L., Tang, K. Y., et al. (2014). Brain-computer interface-based robotic end effector system for wrist and hand rehabilitation: results of a three-armed randomized controlled trial for chronic stroke. Front. Neuroeng. 7, 30. doi:10.3389/fneng.2014.00030.

Bailey, J. O., Bailenson, J. N., and Casasanto, D. (2016). When does virtual embodiment change our minds? Presence 25, 222–233. doi:10.1162/PRES_a_00263.

Banakou, D., Groten, R., and Slater, M. (2013). Illusory ownership of a virtual child body causes overestimation of object sizes and implicit attitude changes. Proc. Natl. Acad. Sci. 110, 12846–12851. doi:10.1073/pnas.1306779110.

Banakou, D., Hanumanthu, P. D., and Slater, M. (2016). Virtual embodiment of white people in a black virtual body leads to a sustained reduction in their implicit racial bias. Front. Hum. Neurosci. 10, 601. doi:10.3389/fnhum.2016.00601.

Biasiucci, A., Leeb, R., Iturrate, I., Perdikis, S., Al-Khodairy, A., Corbet, T., et al. (2018). Brain-actuated functional electrical stimulation elicits lasting arm motor recovery after stroke. Nat. Commun. 9, 2421. doi:10.1038/s41467-018-04673-z.

Carrasco, D. G., and Cantalapiedra, J. A. (2016). Effectiveness of motor imagery or mental practice in functional recovery after stroke: a systematic review. Neurol. (English Ed. 31, 43–52. doi:10.1016/j.nrleng.2013.02.008.

Cincotti, F., Pichiorri, F., Aricò, P., Aloise, F., Leotta, F., De Vico Fallani, F., et al. (2012). EEG-based Brain-Computer Interface to support post-stroke motor rehabilitation of the upper limb. in 2012 Annual International Conference of the IEEE Engineering in Medicine and Biology Society (San Diego, California, USA: IEEE), 4112–4115.

Dechent, P., Merboldt, K.-D., and Frahm, J. (2004). Is the human primary motor cortex involved in motor imagery? Cogn. Brian Res. 19, 138–144. doi:10.1016/j.cogbrainres.2003.11.012.

Delorme, A., and Makeig, S. (2004). EEGLAB: an open source toolbox for analysis of single-trial EEG dynamics including independent component analysis. J. Neurosci. Methods 134, 9–21.

Franceschini, M., Ceravolo, M. G., Agosti, M., Cavallini, P., Bonassi, S., Dall’Armi, V., et al. (2012). Clinical relevance of action observation in upper-limb stroke rehabilitation. Neurorehabil. Neural Repair 26, 456–462. doi:10.1177/1545968311427406.

Frolov, A. A., Mokienko, O., Lyukmanov, R., Biryukova, E., Kotov, S., Turbina, L., et al. (2017). Post-stroke Rehabilitation Training with a Motor-Imagery-Based Brain-Computer Interface (BCI)-Controlled Hand Exoskeleton: A Randomized Controlled Multicenter Trial. Front. Neurosci. 11, 400. doi:10.3389/fnins.2017.00400.

Garrison, K. A., Aziz-Zadeh, L., Wong, S. W., Liew, S.-L., and Winstein, C. J. (2013). Modulating the motor system by action observation after stroke. Stroke 44, 2247–2253. doi:10.1161/STROKEAHA.113.001105.

Garrison, K. A., Winstein, C. J., and Aziz-Zadeh, L. (2010). The mirror neuron system: a neural substrate for methods in stroke rehabilitation. Neurorehabil. Neural Repair 24, 404–412. doi:10.1177/1545968309354536.

Guerra, Z. F., Bellose, L. C., Danielli Coelho De Morais Faria, C., and Lucchetti, G. (2018). The effects of mental practice based on motor imagery for mobility recovery after subacute stroke: Protocol for a randomized controlled trial. Complement. Ther. Clin. Pract. 33, 36–42. doi:10.1016/j.ctcp.2018.08.002.

Jackson, P. L., Lafleur, M. F., Malouin, F., Richards, C. L., and Doyon, J. (2003). Functional cerebral reorganization following motor sequence learning through mental practice with motor imagery. Neuroimage 20, 1171–1180. doi:10.1016/S1053-8119(03)00369-0.

Kennedy, R. S., Lane, N. E., Berbaum, K. S., and Michael, G. (1993). Simulator sickness questionnaire: an enhanced method for quantifying simulator sickness. Int. J. Aviat. Psychol. 3, 203–220. doi:10.1207/s15327108ijap0303_3.

Kilteni, K., Bergstrom, I., and Slater, M. (2013). Drumming in immersive virtual reality: the body shapes the way we play. IEEE Trans. Vis. Comput. Graph. 19, 597–605. doi:10.1109/TVCG.2013.29.

Kilteni, K., Normand, J.-M., Sanchez-Vives, M. V., and Slater, M. (2012). Extending body space in immersive virtual reality: a very long arm illusion. PLoS One 7, e40867. doi:10.1371/journal.pone.0040867.

Leeb, R., Lee, F., Keinrath, C., Scherer, R., Bischof, H., and Pfurtscheller, G. (2007). Brain-computer communication: motivation, aim, and impact of exploring a virtual apartment. IEEE Trans. Neural Syst. Rehabil. Eng. 15, 473–482. doi:10.1109/TNSRE.2007.906956.

Mcmahon, M., and Schukat, M. (2018). A low-Cost, Open-Source, BCI-VR Game Control Development Environment Prototype for Game based Neurorehabilitation. in IEEE Games, Entertainment, Media Conference (GEM), 1–9.

Naito, E., Kochiyama, T., Kitada, R., Nakamura, S., Matsumura, M., Yonekura, Y., et al. (2002). Internally simulated movement sensations during motor imagery activate cortical motor areas and the cerebellum. J. Neurosci. 22, 3683–3691. doi:20026282.

Ono, T., Shindo, K., Kawashima, K., Ota, N., Ito, M., Ota, T., et al. (2014). Brain-computer interface with somatosensory feedback improves functional recovery from severe hemiplegia due to chronic stroke. Front. Neuroeng. 7. doi:10.3389/fneng.2014.00019.

Osimo, S. A., Pizarro, R., Spanlang, B., and Slater, M. (2015). Conversations between self and self as Sigmund Freud—A virtual body ownership paradigm for self counselling. Sci. Rep. 5, 13899. doi:10.1038/srep13899.

Pavone, E. F., Tieri, G., Rizza, G., Tidoni, E., Grisoni, L., and Aglioti, S. M. (2016). Embodying others in immersive virtual reality: electro-cortical signatures of monitoring the errors in the actions of an avatar seen from a first-person perspective. J. Neurosci. 36, 268–279. doi:10.3389/fpsyg.2016.01260.

Pichiorri, F., Morone, G., Petti, M., Toppi, J., Pisotta, I., Molinari, M., et al. (2015). Brain-computer interface boosts motor imagery practice during stroke recovery. Ann. Neurol. 77, 851–865. doi:10.1002/ana.24390.

Prochnow, D., Bermúdez i Badia, S., Schmidt, J., Duff, A., Brunheim, S., Kleiser, R., et al. (2013). A functional magnetic resonance imaging study of visuomotor processing in a virtual reality-based paradigm: Rehabilitation Gaming System. Eur. J. Neurosci. 37, 1441–1447. doi: 10.1111/ejn.12157.

Ramos-Murguialday, A., Broetz, D., Rea, M., Läer, L., Yilmaz, Ö., Brasil, F. L., et al. (2013). Brain-machine interface in chronic stroke rehabilitation: a controlled study. Ann. Neurol. 74, 100–108. doi:10.1002/ana.23879.

Slater, M., and Sanchez-Vives, M. V. (2016). Enhancing our lives with immersive virtual reality. Front. Robot. AI 3, 74. doi:10.3389/frobt.2016.00074.

Spicer, R., Anglin, J., Krum, D. M., and Liew, S. L. (2017). REINVENT: A low-cost, virtual reality brain-computer interface for severe stroke upper limb motor recovery. in IEEE Virtual Reality (IEEE), 385–386. doi:10.1109/VR.2017.7892338.

Tung, S. W., Guan, C., Ang, K. K., Phua, K. S., Wang, C., Zhao, L., et al. (2013). Motor imagery BCI for upper limb stroke rehabilitation: an evaluation of the EEG recordings using coherence analysis. in 2013 35th Annual International Conference of the IEEE Engineering in Medicine and Biology Society (EMBC) (IEEE), 261–261.

Uktveris, T., and Jusas, V. (2018). Development of a modular board for eeg signal acquisition. Sensors (Switzerland) 18, 2140. doi:10.3390/s18072140.

Vecchiato, G., Tieri, G., Jelic, A., De Matteis, F., Maglione, A. G., and Babiloni, F. (2015). Electroencephalographic Correlates of Sensorimotor Integration and Embodiment during the Appreciation of Virtual Architectural Environments. Front. Psychol. 6, 1944. doi:10.3389/fpsyg.2015.01944.

Vourvopoulos, A., and Bermúdez i Badia, S. (2016). Motor priming in virtual reality can augment motor-imagery training efficacy in restorative brain-computer interaction: a within-subject analysis. J. Neuroeng. Rehabil. 13, 69. doi:10.1186/s12984-016-0173-2.

Welch, P. D. (1967). The Use of Fast Fourier Transform for the Estimation of Power Spectra: A Method Based on Time Averaging Over Short, Modified Periodograms. IEEE Trans. Audio Electroacoust. 15, 70–73.

Witmer, B. G., and Singer, M. J. (1998). Measuring presence in virtual environments: a presence questionnaire. Presence 7, 225–240. doi:10.1162/105474698565686.

Yee, N., and Bailenson, J. N. (2007). The Proteus effect: The effect of transformed self-representation on behavior. Hum. Commun. Res. 33, 271–290. doi:10.1111/j.1468-2958.2007.00299.x.

Zhu, M.-H., Wang, J., Gu, X.-D., Shi, M.-F., Zeng, M., Wang, C.-Y., et al. (2015). Effect of action observation therapy on daily activities and motor recovery in stroke patients. Int. J. Nurs. Sci. 2, 279–282. doi:10.1016/j.ijnss.2015.08.006.

