## Supplemental Figures for "Embodiment is related to better performance on an immersive brain computer interface in head-mounted virtual reality: A pilot study"

**Supplemental Figure 1. Individual participant EEG activity for C3. (A)** Analysis of the relative PSD to baseline of C3 for the alpha band at the group-level (left) and at the individual-level (right). For individual subjects, for relative alpha levels during the task, there were significant differences between Screen and HMD-VR in nine participants. From those, three participants had significantly lower alpha (greater desynchronization) during the Screen condition and six participants had significantly lower alpha (greater desynchronization) during the HMD-VR condition. **(B)** Analysis of the relative PSD to baseline of C3 for the beta band at the group-level (left) and at the individual-level (right). For individual subjects, for relative beta levels during the task, there were significant differences between Screen and HMD-VR in ten participants. From those, five participants had significantly lower beta during the Screen condition and five participants had significantly lower beta during the HMD-VR condition.

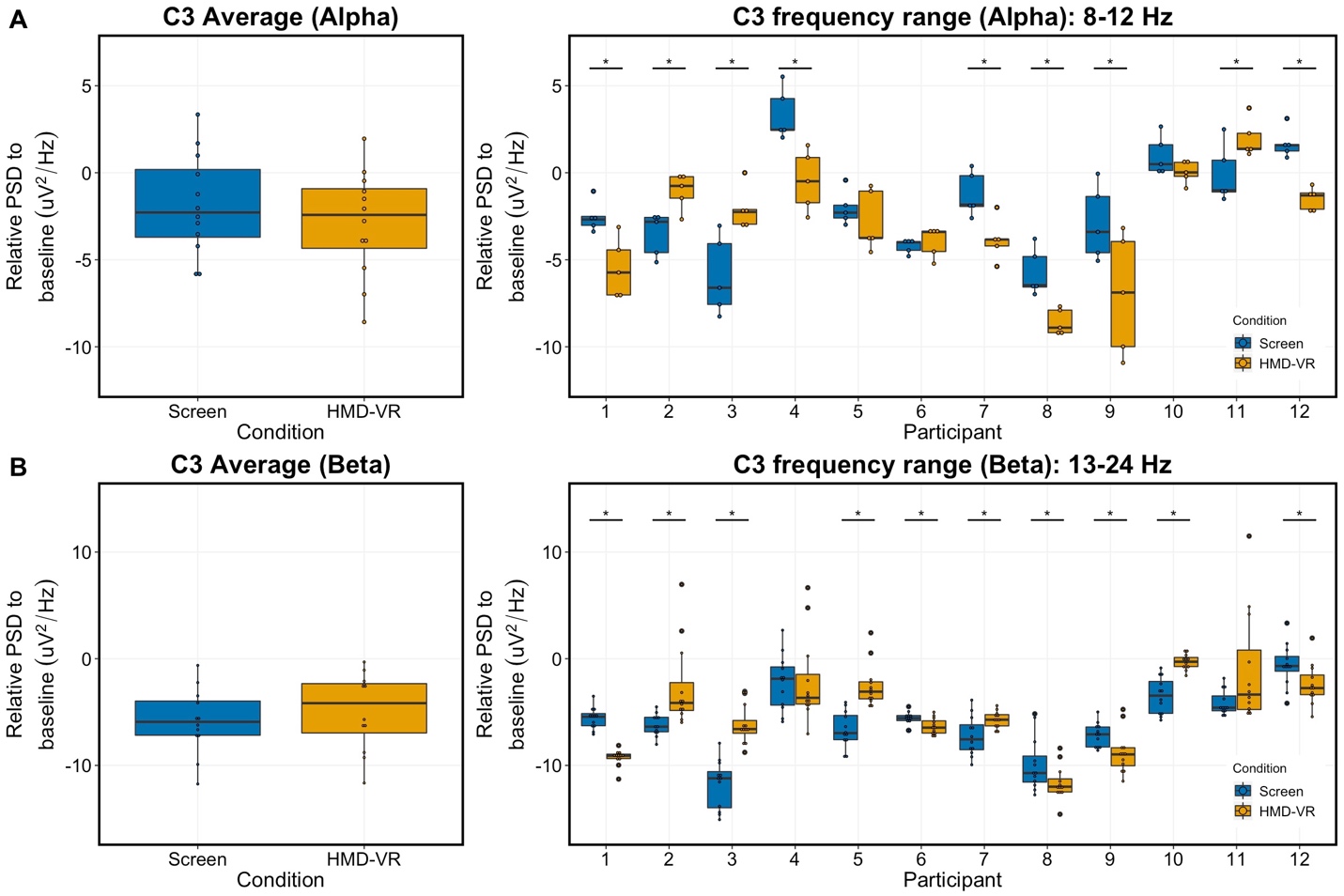

**Supplemental Figure 2. Individual participant EEG activity for C4. (A)** Analysis of the relative PSD to baseline of C4 for the alpha band at the group-level (left) and at the individual-level (right). There were significant differences between Screen and HMD-VR in eleven participants. From those, four participants had significantly lower alpha during the Screen condition and seven participants had significantly lower alpha during the HMD-VR condition. **(B)** Analysis of the relative PSD to baseline of C4 for the beta band at the group-level (left) and at the individual-level (right). There were significant differences in the beta band between Screen and HMD-VR in eleven participants. From those, five participants had significantly lower beta during the Screen condition and six participants had significantly lower beta during the HMD-VR condition.

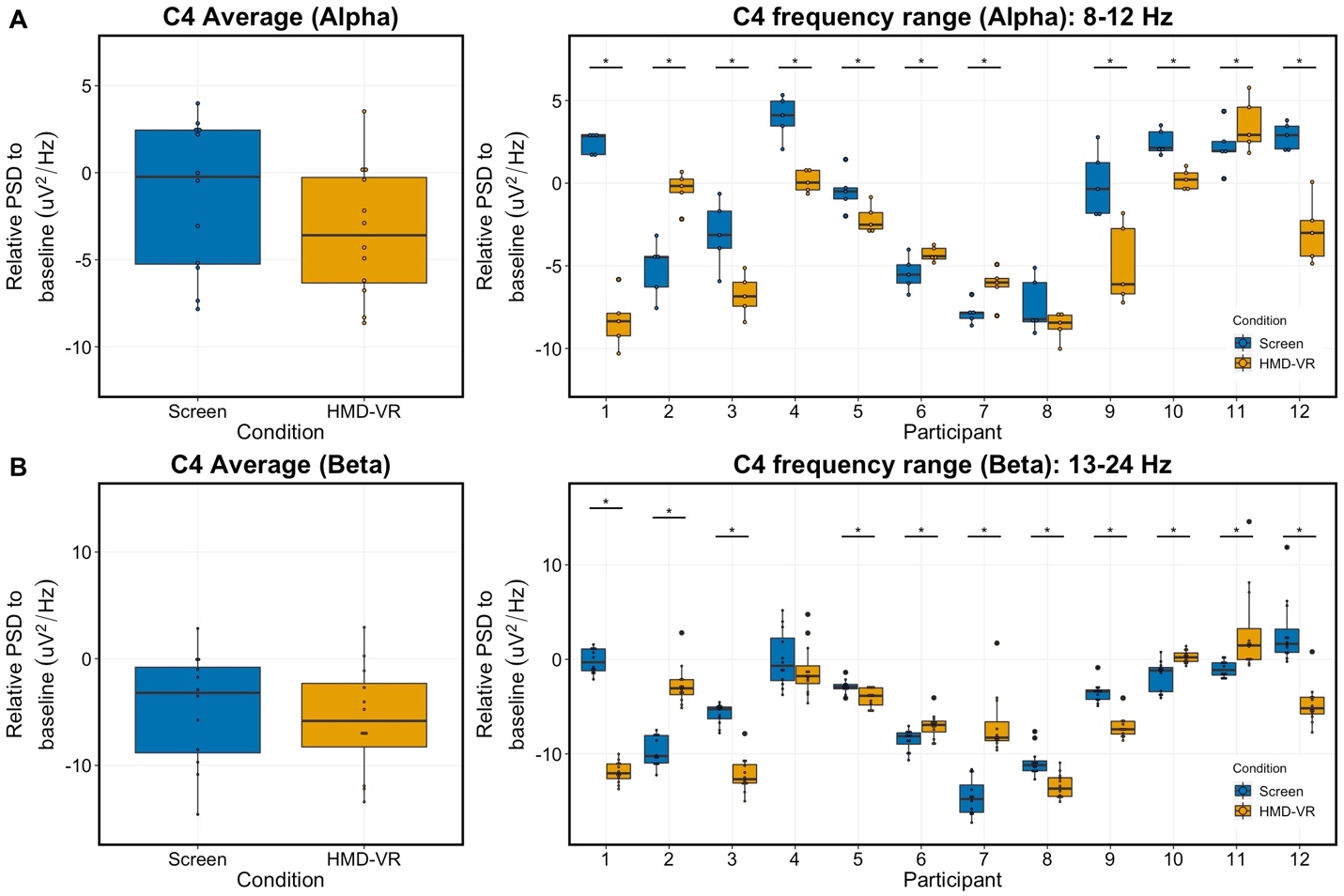
